# Lack of synergy between AR targeted therapies and PARP inhibitors in homologous recombination-proficient prostate cancer

**DOI:** 10.1101/2025.06.02.657429

**Authors:** Nicole A. Traphagen, Esmé Wheeler, Rong Li, Feng Lu, Buraq Ahmed, Alok K. Tewari, Steven P. Balk, Peter S. Nelson, Eva Corey, Henry Long, Alan D. D’Andrea, Xintao Qiu, Myles Brown

## Abstract

Recent clinical trials have explored the combination of androgen receptor (AR) pathway inhibitors and poly (ADP-ribose) polymerase (PARP) inhibitors as a potential treatment for castration-resistant prostate cancer. This combination treatment is based on the premise that AR directly regulates expression of DNA repair genes, leading to synergy between PARP and AR inhibition. Despite some promising preclinical evidence, this combination therapy has shown limited efficacy in patients with homologous recombination (HR)-proficient tumors. To investigate this discrepancy between preclinical and clinical results, we profiled the effects of PARP inhibition in prostate cancer models in the presence or absence of AR inhibition. Surprisingly, AR inhibition impaired response to PARP inhibitors in castration-sensitive cells and had no effect on response in castration-resistant cells. AR inhibition also did not regulate DNA repair in either the castration-resistant or castration-sensitive setting. Instead, we find that cell cycle progression is required for response to PARP inhibition in homologous-recombination proficient prostate cancer.

**STATEMENT OF SIGNIFICANCE:** Androgen deprivation does not inhibit DNA repair and does not synergize with PARP inhibition in prostate cancer with intact homologous recombination repair.

## INTRODUCTION

Inhibitors of poly (ADP-ribose) polymerase (PARP) are synthetically lethal with loss of the homologous recombination (HR) DNA repair pathway. PARP inhibitors (PARPi) are approved for use in advanced prostate cancer in HR-deficient tumors, including tumors with germline or somatic alterations in *BRCA1* or *BRCA2*. However, whether there is a role for PARPi in HR-proficient tumors remains an open question.

Inhibition of the androgen receptor (AR) signaling through androgen deprivation therapy or treatment with AR antagonists is the standard of care for prostate cancer, which is typically dependent upon AR in both early stage (castration-sensitive) and late stage (castration-resistant) disease. Prior studies have shown that the AR regulates expression of DNA damage response genes, including *XRCC2, XRCC3, and PRKDC* (1), the HR genes *BRCA1* and *BRCA2* (2), and *PARP1* (3). This model of AR regulation of the DNA damage response suggests that AR signaling inhibition suppresses expression of DNA damage response genes, limiting DNA repair and leading to synergy with DNA damaging agents such as ionizing radiation and PARPi. However, clinical studies assessing the efficacy of combination AR inhibition and PARPi have had mixed results (4–6). In these studies, most patients who benefitted from the combination treatment had HR-deficient tumors, and patients with loss of *BRCA1/2* had the greatest improvement in response (4–6). Patients with HR-proficient tumors exhibited only a small improvement in progression-free survival and overall survival (4, 5) or no improvement in progression-free survival (6) upon the addition of PARPi to AR signaling inhibition, compared to AR signaling inhibition alone. While several of these studies showed PARPi treatment resulted in improved outcomes in HR-proficient patients, the improvement in progression free survival was more modest than would be expected if AR signaling inhibition and PARPi truly synergized in this context. While these trials led to approval of combination talazoparib (PARPi) and enzalutamide (AR inhibitor) in patients with metastatic castration resistant prostate cancer (CRPC) with or without HR defects in the European Union, the U.S. Food and Drug Administration recently declined to grant approval for talazoparib in combination with enzalutamide treatment for patients with HR-proficient prostate cancer (7).

There are 17 PARP family enzymes with diverse roles in DNA repair, transcription, and stress responses. Most PARPi, including those FDA approved for use in prostate cancer (i.e. olaparib, rucaparib, and talazoparib) inhibit both PARP1 and PARP2, although the PARP1-selective inhibitor AZD-5305 (saruparib) has recently entered clinical trials. PARP1 and PARP2 have redundant and distinct roles in repair of single-stranded DNA breaks (8). PARP1 has been shown to remodel chromatin and mediate transcription through diverse mechanisms (9–12). PARP2 further plays a specific role in enhancing AR transcriptional activity through FOXA1 mediated effects (13). Given the roles of PARP1 and PARP2 in transcription, and the role of AR in regulating the DNA damage response, we set out to untangle the complex interplay between AR, the DNA damage response, and PARP inhibition in prostate cancer.

## RESULTS

### AR inhibition does not synergize with PARP inhibition in castration-sensitive or castration-resistant models of prostate cancer

We first set out to confirm that AR inhibition and PARP inhibition synergize in preclinical prostate cancer models. For these experiments we chose to use the PARP1/2 inhibitors olaparib and rucaparib, which are FDA-approved for HR-deficient CRPC, as well as the PARP1-selective inhibitor AZD-5305 (saruparib). To our surprise, rather than synergizing with PARP inhibition, androgen deprivation strongly inhibited response to all three PARPi tested in castration-sensitive prostate cancer (CSPC) LNCaP cells (Fig. 1A).

**Figure 1.**
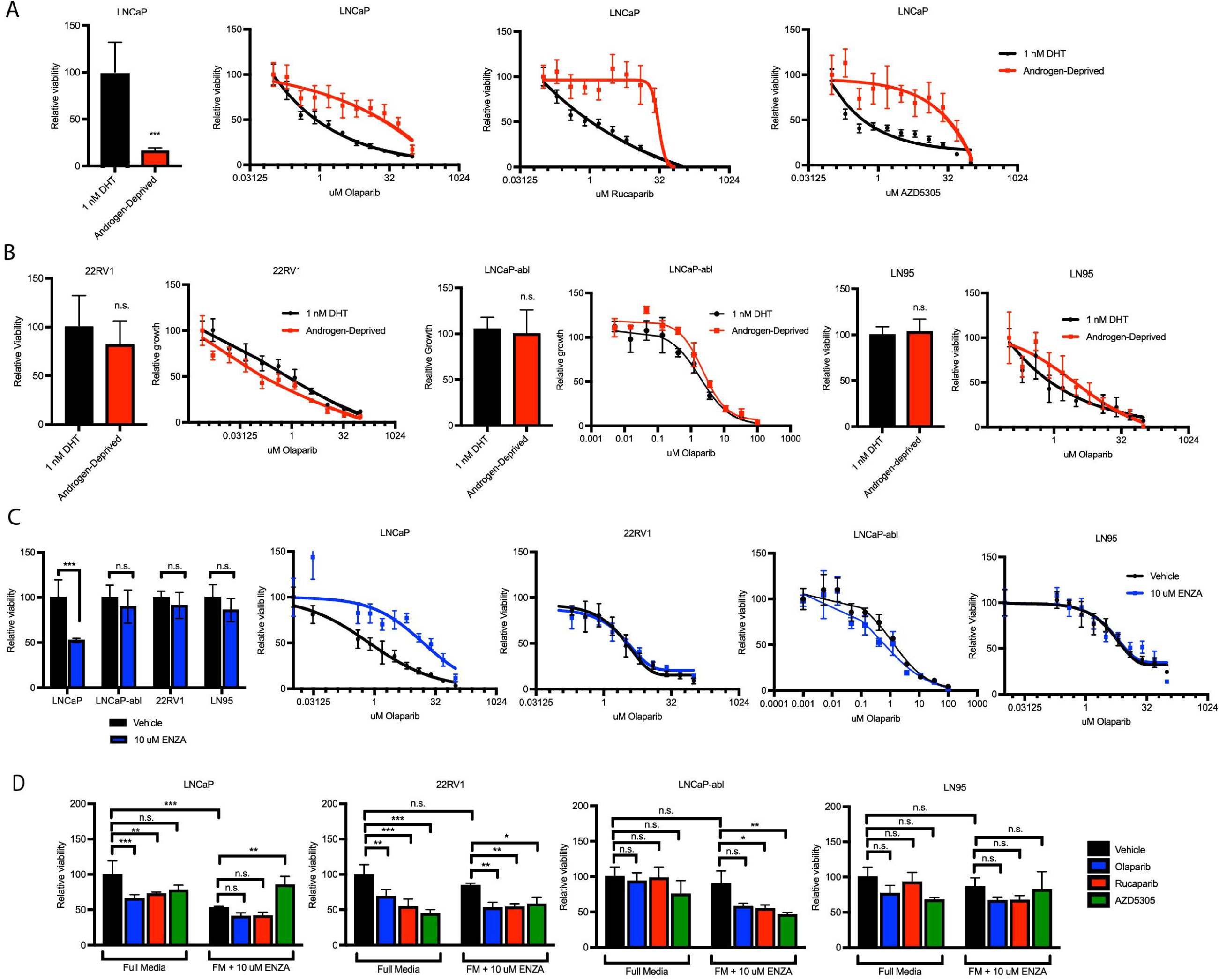
AR inhibition does not alter sensitivity to PARP inhibition in castration-resistant prostate cancer. (A) LNCaP cells were seeded in androgen-deprived media ± 1 nM DHT and treated in quadruplicate as indicated for 7 d. Viability was assessed using Cell Titer Glo. (B) Castration-resistant cell lines were seeded in androgen-deprived media ± 1 nM DHT and treated as indicated for 7 d. Viability was assessed using Cell Titer Glo. (C) Cells were seeded in growth media ± 10 μM enzalutamide (ENZA) and treated as indicated for 7 d. Viability was assessed using Cell Titer Glo. (D) Cells were seeded in growth media and treated ± 1 μM PARPi and ± 10 μM ENZA as indicated for 7 d. Viability was assessed using Cell Titer Glo. Data are shown as mean ± SD. *p<0.05, **p<0.005, ***p<0.005, n.s. = not significant by Student’s two-tailed t-test (A-C) or one-way ANOVA with Bonferroni correction for multiple comparisons (D).

Combination PARPi and AR inhibition is being tested clinically in the setting of CRPC, so we expanded our studies to include the CRPC models 22RV1, LN95, and LNCaP-abl. In these castration-resistant cells, which are not growth inhibited by androgen-deprivation, androgen deprivation had no effect on response to PARPi (Fig. 1B). We saw similar results upon treatment with the AR antagonist enzalutamide. Enzalutamide treatment greatly increased the IC50 of PARPi in CSPC LNCaP cells but had no effect on response to PARPi in CRPC models (Fig 1C). At clinically relevant concentrations (10 uM enzalutamide and 1 uM PARPi) combination treatment with enzalutamide and PARP inhibition did not confer any added benefit compared to either drug alone (Fig. 1D). While unexpected, these results strongly suggest that AR inhibition does not synergize with PARPi in HR-proficient prostate cancer.

### Treatment with PARP inhibitors modulates expression of androgen responsive genes

To better understand the effects of PARP inhibition in the presence or absence of AR inhibition, we performed RNA-sequencing of PARPi treated cells. For these experiments, we chose to use castration-sensitive LNCaP and castration-resistant 22RV1 cells. We first confirmed that treatment with PARP inhibitors had no effect on AR expression (Fig. 2A). Cells were then treated with PARPi in either androgen-deprived (vehicle) or DHT-stimulated conditions (Fig 2B). RNA-sequencing indicated that treatment with DHT induced a transcriptional response in both 22RV1 and LNCaP cells at 4 h and 16 h of treatment (Fig. S1A), which corresponded with enrichment for the Hallmark Androgen Response gene set at these timepoints (Fig. S1B).

**Figure 2.**
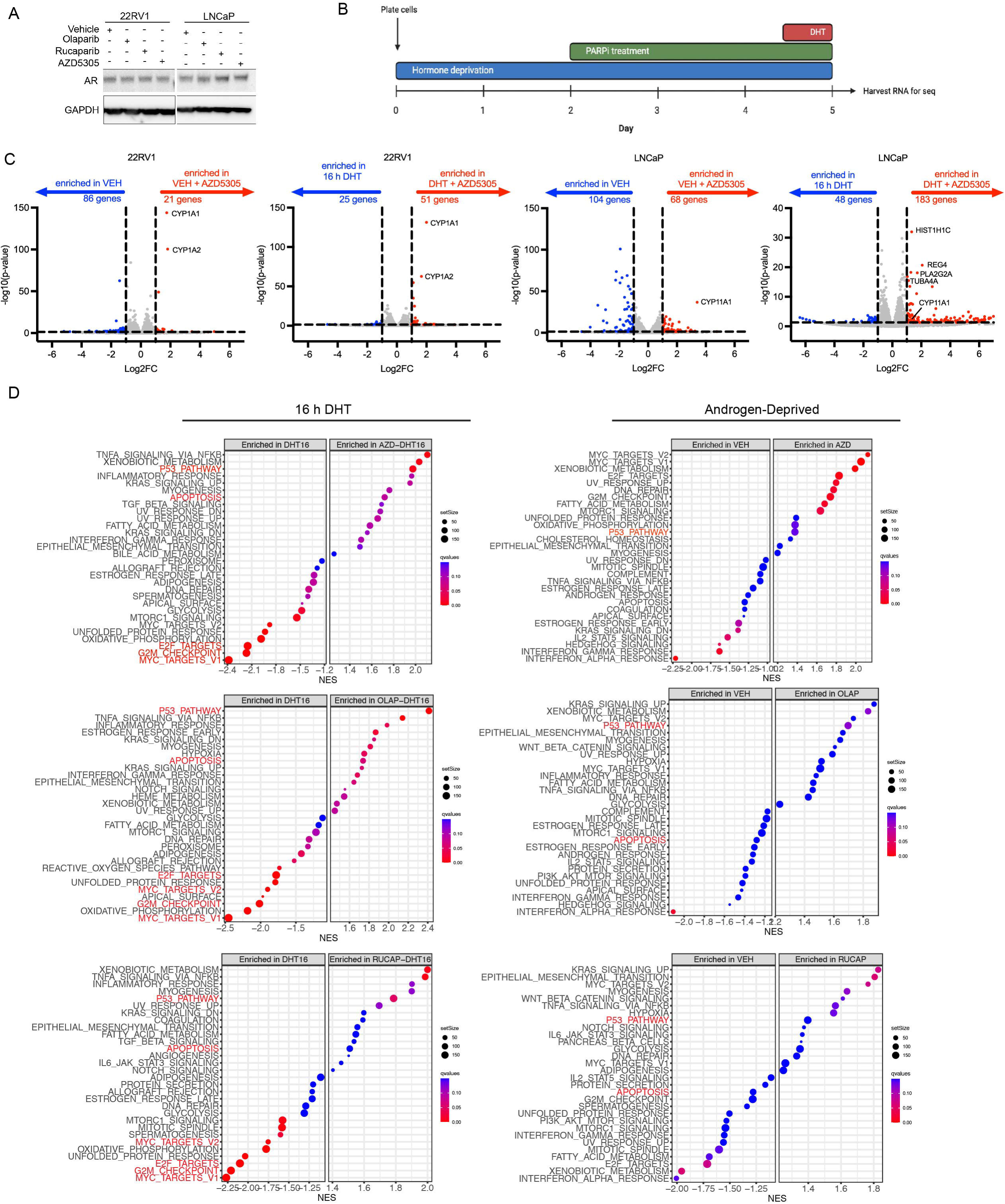
PARP inhibition induces changes in cell cycle and DNA damage response pathways. (A) Cells were seeded in growth media and treated with 5 μM PARPi for 3 d. Cells were lysed and protein harvested for immunoblot. (B) Experimental design for RNA-sequencing experiments. Cells were treated ± 5 μM PARPi and ± 1 nM DHT. (C) Genes with significantly differential expression (Log2FC >|1|, p-adjusted <0.05) upon treatment with AZD-5305 in the presence or absence of DHT stimulation. (D) Normalized enrichment scores for Hallmarks pathways in cells treated with PARPi in the presence or absence of DHT.

All three PARPi strongly induced expression of CYP genes in our experiments, including *CYP1A* and *CYP1B* (Fig. 2C, Fig. S2-S3). However, no other genes were consistently altered upon PARPi treatment. To gain a better understanding of the effects of PARPi treatment on cells under androgen-deprived or DHT stimulated conditions, we next looked at enrichment scores for Hallmark pathways (Fig. 2D). The p53 pathway was enriched upon treatment with all three PARPi in both DHT-stimulated and androgen-deprived conditions; however, it was more strongly enriched in cells stimulated with DHT. Cell cycle-related pathways (E2F targets, G2/M Checkpoint, MYC targets) were decreased in PARPi treated cells upon DHT-stimulation but not in androgen-deprived conditions. Similarly, apoptosis was enriched upon PARPi treatment only in DHT-stimulated cells. This suggests that PARPi treatment elicits growth-inhibitory effects in DHT-treated cells but not androgen-deprived cells, consistent with our growth data in Figure 1.

We next investigated the effect of PARPi specifically on AR-regulated genes, using all genes that were significantly altered at either 4 h or 16 h of DHT treatment. In 22RV1 cells PARPi treatment inhibited DHT-induced expression and suppressed DHT-induced repression of a subset of androgen regulated genes (22RV1 Clusters 2 and 4, Fig. 3A). Genes in these clusters corresponded with The Hallmark Androgen Response gene set, as well as cell cycle and metabolic gene sets (Fig. S4A). In LNCaP cells, PARPi treatment repressed DHT-induced expression of only a small subset of genes (LNCaP Cluster 1, Fig. 3A). GSEA analysis of this gene sets did not reveal any specific pathway enriched in this cluster of genes. (Fig. S4B). Interestingly, there was no apparent difference in AR-driven gene expression upon PARP1 inhibition compared to PARP1/2 inhibition (Fig. 3A), despite the distinct proposed role of PARP2 in AR-driven transcription (13).

**Figure 3.**
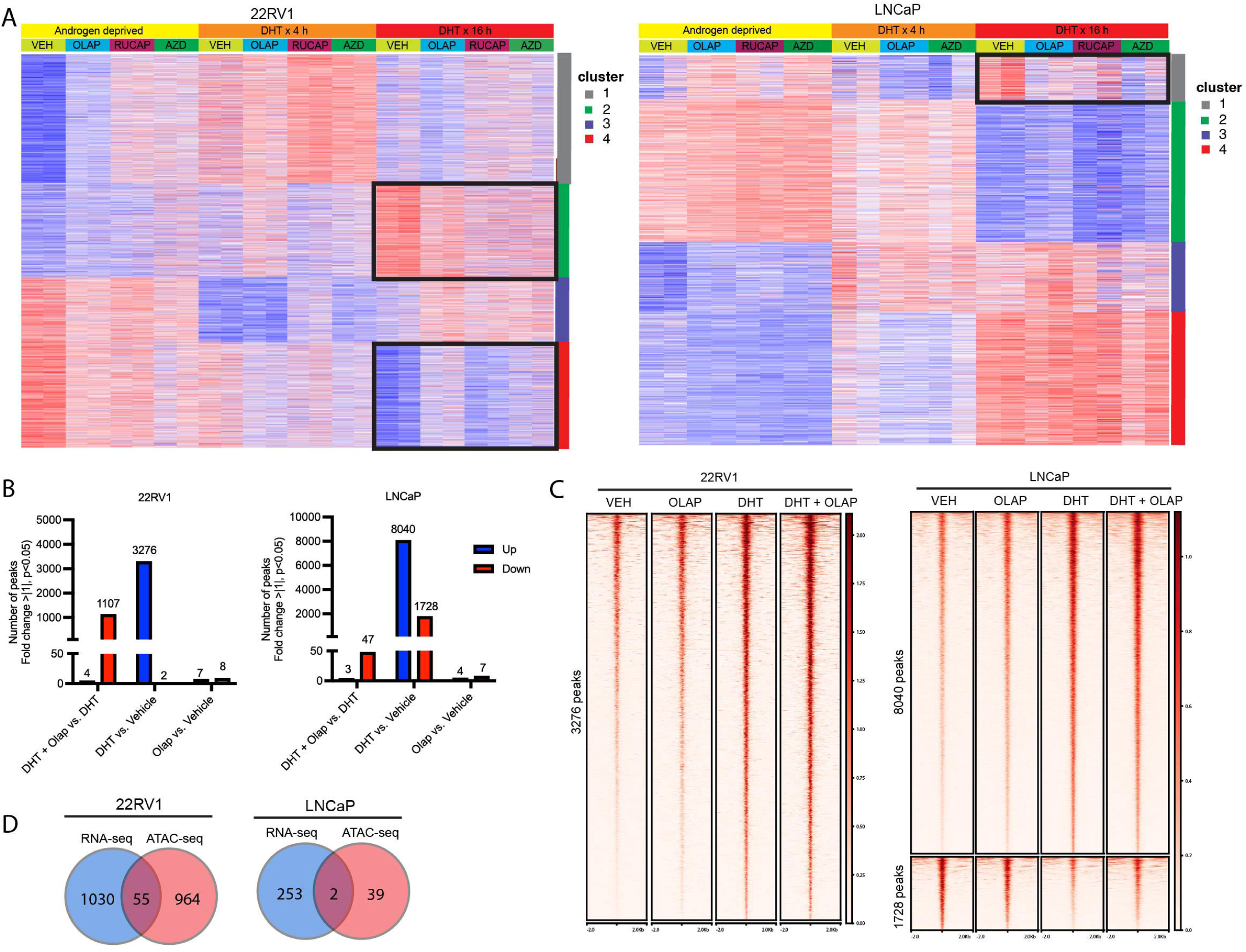
PARP inhibition modulates AR-driven transcription. (A) All genes with significantly differential expression (p-adjusted <0.05) upon treatment with DHT are shown. Boxes indicate clusters of genes with induced or repressed expression upon DHT stimulation that are altered by PARPi treatment. (B) ATAC-seq was performed on cells treated similarly to 2B. Cells were androgen-deprived for two days, treated ± 5 μM olaparib for 3 d, and treated ± 1 nM DHT for the final 24 h prior to harvesting nuclei for ATAC-seq. The number of significantly differential peaks (Log2FC >|1|, p-adjusted <0.05) for each condition is shown. (C) Tornado plot of all sites with significantly different accessibility upon treatment with DHT, shown for the four treatment conditions. (D) Venn diagram showing overlap between AR-regulated genes with expression modulated by PARPi (as shown in Fig. 2A; 22RV1 Clusters 2 and 4, LNCaP Cluster 1) and AR-regulated ATAC-seq peaks with accessibility modulated by PARPi (as shown in Fig. 2B; DHT+ OLAP vs. DHT).

PARP1/2 can modulate chromatin structure but can also more directly regulate transcription (14–16). We performed ATAC-seq on cells treated with olaparib in the presence or absence of DHT stimulation to determine whether the observed changes in AR-driven transcription were regulated at the chromatin or transcriptional level. DHT treatment induced changes in chromatin accessibility following 24h of DHT treatment (Fig. 3B and S5A). In both LNCaP and 22RV1 cells PARPi treatment had little effect on chromatin accessibility under androgen-deprived conditions (Fig. 3B-C). In both cell lines PARPi treatment resulted in decreased chromatin accessibility at a subset of sites (Fig. 3B). Consistent with the RNA-sequencing data, a larger proportion of differentially accessible sites were found in 22RV1 cells compared to LNCaP cells (1107 peaks in 222RV1 vs. 47 peaks in LNCaP; Fig 3B).

To determine whether these changes in chromatin accessibility drive the changes in AR-driven gene expression observed upon PARP inhibitor treatment we compared the genes nearby differential ATAC-seq peaks with genes that are downregulated upon PARPi treatment. In both cell lines we found little overlap between these two gene sets (Fig. 3D), indicating that PARPi-induced changes in gene expression are not due to changes in chromatin accessibility at these sites. As an example, *LRRC4,* a gene with induced expression by DHT and regulated by PARPi, shows increasing accessibility with DHT treatment but is unchanged by olaparib treatment (Fig. S5B). This further supports the conclusion that PARPi-induced changes in gene expression are regulated at the transcriptional level.

### Androgen deprivation or AR inhibition does not repress DNA repair

Prior studies have reported that AR inhibition with the direct AR antagonist enzalutamide inhibits expression of DNA repair genes, including *BRCA2*, leading to synergy with PARP inhibition (2). Since we did not observe synergy between PARPi and AR inhibition in these models, we investigated the effects of AR inhibition on DNA repair genes in our sequencing data. In both 22RV1 and LNCaP cells, androgen deprivation downregulated expression of Hallmark Androgen Response genes (Fig. 4A). Androgen deprivation had no effect on expression of DNA damage response genes in 22RV1 cells. Surprisingly, we observed a modest increase in expression of DNA repair genes in LNCaP cells, in direct contrast to previously published reports suggesting that androgen deprivation decreases expression of these DNA repair genes (1, 2, 17). Our ATAC-seq data also showed that androgen deprivation had no effect on the accessibility of *BRCA1* and *BRCA2* genes (Fig. 4B). Contrary to prior reports, these data suggest that AR does not play a direct role in regulating DNA repair capacity.

**Figure 4.**
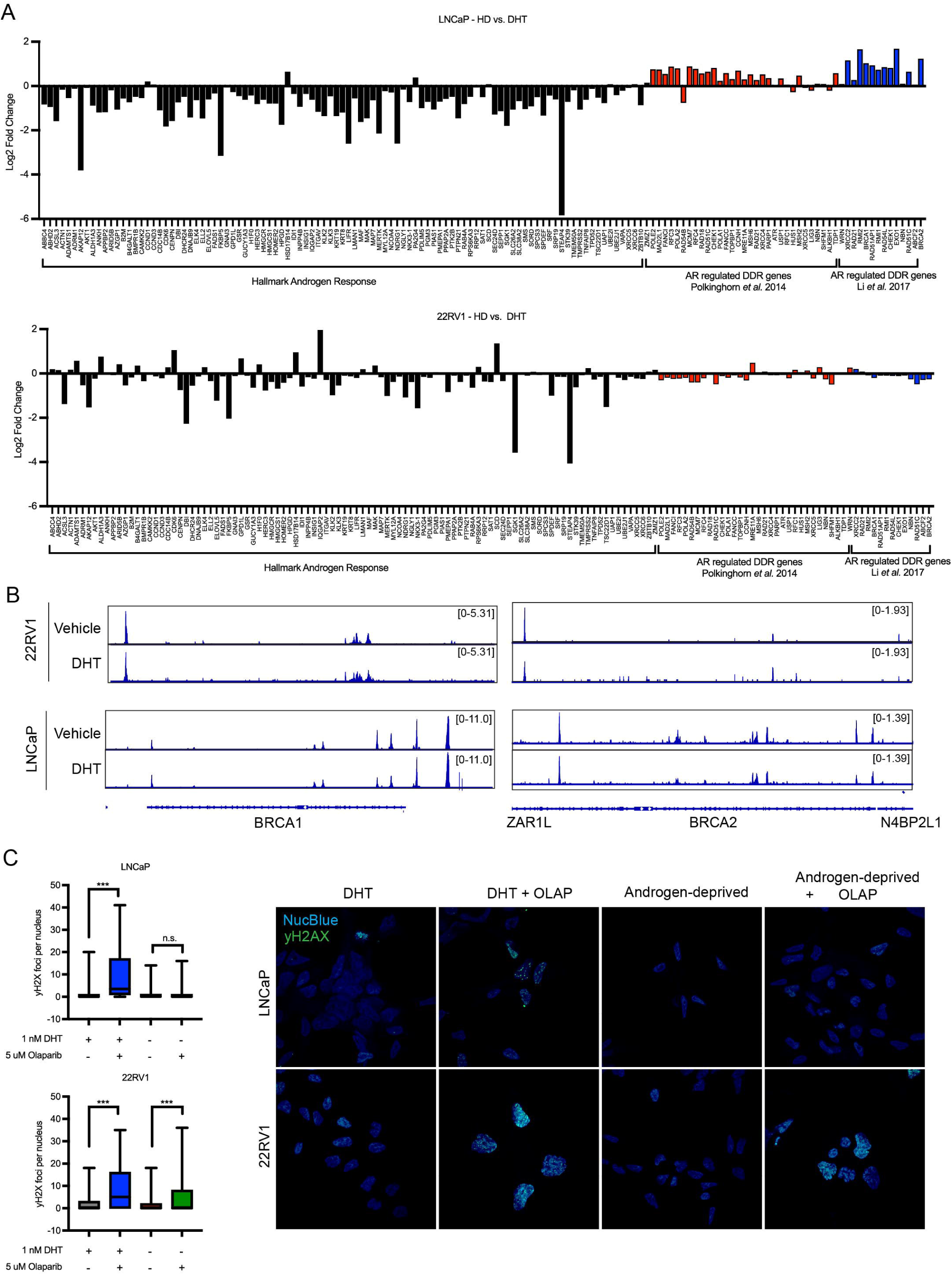
AR does not directly regulate expression of DNA repair genes. (A) Log2FC of genes in the Hallmark Androgen Response pathway and DNA repair genes previously associated with AR regulation (Refs. 2 and 16) in vehicle treated vs. 16 h DHT treated samples. (B) Chromatin accessibility peaks from ATAC-seq data in vehicle treated vs. 24 h DHT treated samples. (C) Cells were seeded in androgen deprived media and treated ± 1 nM DHT, treated ± 5 μM olaparib for 72 h. DNA damage was assessed by staining for nuclear γH2AX foci, and staining quantified in >100 cells per condition. *p<0.05, **p<0.005, ***p<0.005, n.s. = not significant by one-way ANOVA with Bonferroni correction for multiple comparisons.

To functionally assess DNA damage induced by PARPi in combination with AR signaling inhibition, we treated cells with olaparib in the presence or absence of DHT for 72 h and assessed DNA damage through γH2AX foci formation. In LNCaP cells, androgen deprivation suppressed induction of DNA damage upon olaparib treatment. In 22RV1 cells, DNA damage was observed regardless of whether cells were cultured in the presence or absence of DHT. We observed similar results in cells treated with enzalutamide and olaparib (Fig. S6).

Together, these data indicate that AR pathway inhibition does not inhibit the DNA damage response in prostate cancer cells or increase PARPi-induced DNA damage in HR-proficient cells.

### Cell cycle progression is required for response to PARP inhibition

While continuous treatment with AR signaling inhibition is the standard treatment for prostate cancer, interrupting androgen deprivation through treatment with supraphysiological androgens (termed ‘bipolar androgen therapy; BAT) can improve response to ADT in CRPC (18–21). Our finding that HR-proficient cells have improved response to PARP inhibitors in the presence of physiological levels of androgens (Fig. 1A) raises the question of whether PARP inhibition would be more effective in the context of supraphysiological androgen levels rather than ADT. Others have found that treatment with olaparib enhances the anti-cancer effects of supraphysiological androgens and BAT regardless of HR mutation status (22, 23). Similar results have been reported in advanced breast cancer, where olaparib treatment synergized with estrogen treatment regardless of HR mutation status (24). We used LNCaP cells to test this hypothesis, as these cells proliferate upon treatment with 1 nM DHT but are growth inhibited by treatment with 10 nM DHT (Fig. 5A). Treatment with 10 nM DHT strongly inhibited response to olaparib in LNCaP cells (Fig. 5B), to an even greater extent than androgen-deprivation. This suggests that cell cycle effects, and not AR transcriptional effects, underlie the difference in response to PARP inhibitors upon androgen deprivation. This is supported by our finding that 22RV1 and LNCaP cells exhibit changes in AR-driven transcription upon DHT treatment (Fig S1A-B), but only LNCaP cells are growth inhibited and exhibit a suppressed response to PARPi in androgen-deprived media (Fig.1A-B). We therefore reasoned that the decreased response to PARPi in castration-sensitive cells was due to cell cycle changes induced by AR inhibition rather than effects on AR-driven transcription.

**Figure 5.**
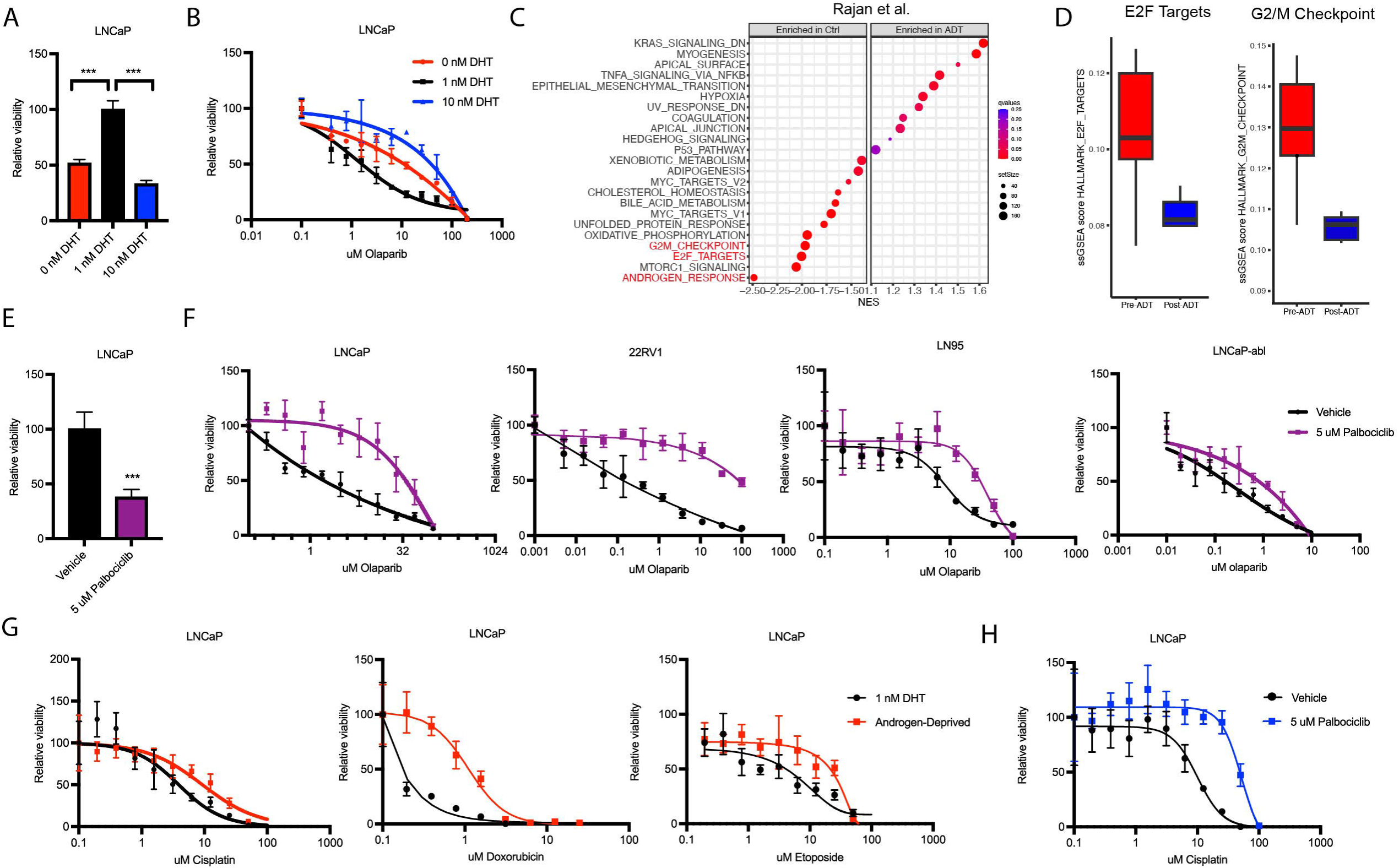
Halting cell cycle progression impairs therapeutic response to PARPi. (A/B) LNCaP cells were seeded in androgen-deprived media and treated with DHT and/or olaparib as indicated in quadruplicate for 7 d prior to assessing cell viability. (C) Normalized enrichment score for Hallmarks pathways in RNA-sequencing data from pre-ADT and post-ADT treated patient samples as described in Ref. 24. (C) Single-sample GSEA scores of cell cycle related Hallmarks pathways in matched pre-ADT and post-ADT samples. (E) LNCaP cells were seeded in full media and treated ± 5 μM palbociclib in quadruplicate for 7 d prior to assessing cell viability. (F) Cells were seeded in growth media ± 5 μM palbociclib and treated with olaparib as indicated in quadruplicate for 7 d prior to assessing cell viability. (G) Cells were seeded in androgen-deprived media ± 1 nM DHT and treated as indicated with DNA damaging chemotherapies for 7 d prior to assessing cell viability. (H) Cells were seeded in full media ± 5 μM palbociclib and treated with cisplatin as indicated for 7 d. Data are shown as mean ± SD, ***p<0.005. by one-way ANOVA with Bonferroni correction for multiple comparisons (A) or Student’s two-tailed t-test (E).

To confirm that androgen deprivation alters cell cycle progression in patients, we analyzed publicly available RNA-sequencing data of matched pre-ADT and post-ADT prostate cancer (25). In this dataset treatment with ADT strongly inhibits gene sets associated with cell cycle progression, including downregulation of Hallmark E2F targets and G2M checkpoint pathways, as well as the Hallmark Androgen Response (Fig. 5C). In patient-matched pre-ADT and post-ADT samples, E2F targets and the G2/M checkpoint gene sets were strongly suppressed following treatment with ADT (Fig. 5D), indicating that ADT alters cell cycle progression in patients.

To more directly test the hypothesis that inhibition of cell cycle impairs response to PARP inhibitors, we first determined the IC50 of the CDK4/6 inhibitor palbociclib in prostate cancer cell lines (Fig. S7A). Treatment with 5 μM palbociclib inhibited growth in all cell lines (Fig. 5E, Fig S7B) and suppressed response to olaparib in both castration-sensitive cells and castration-resistant cells (Fig. 5F). Similar results were observed when palbociclib was combined with rucaparib or AZD-5305 (Fig. S7C).

The primary mechanism of action of PARP inhibitors is induction of DNA damage, and many DNA-damaging therapies have cell-cycle specific effects. We therefore hypothesized that the differences in response observed in the presence or absence of AR signaling inhibition would not be specific to PARPi but extended to all DNA damaging chemotherapies. Indeed, inhibition of AR through androgen deprivation or enzalutamide treatment impaired response to the DNA damaging cytotoxic chemotherapies cisplatin, etoposide, and doxorubicin in LNCaP (Fig. 5G) but not CRPC cells (Fig. S8A-B). Similar to results observed with PARPi, halting cell cycle progression through inhibition of CDK4/6 also inhibited response to these DNA damaging chemotherapies (Fig. 5H, S8C-D). Together, these data indicate that cell cycle progression is required for response to PARPi in HR-proficient prostate cancer. These effects were not unique to PARP inhibitors, but generalizable to all DNA-damaging treatments tested.

## DISCUSSION

In this study we find that PARP inhibitors modulate AR-driven gene expression programs at the transcriptional level, in accordance with prior reports suggesting that PARP1 and PARP2 have roles in mediating transcription (10–13). In our models, inhibition of AR did not regulate DNA damage repair, which likely explains our finding that androgen deprivation does not synergize with PARP inhibition. Instead, we report that interrupting cell cycle progression through either AR inhibition or CDK4/6 inhibition impairs response to PARP inhibition in HR-proficient prostate cancer models.

Interestingly, there were no apparent differences between the effects of PARP1/2 inhibition and PARP1-selective inhibition in our HR-proficient models, despite the proposed unique role of PARP2 in AR-driven transcription (13). PARP1-selective inhibitors, which are currently in early-phase clinical testing, were developed based on the observation that the myelosuppressive toxicity of PARP1/2 inhibitors is primarily due to effects on PARP2, while anticancer effects are primarily due to effects on PARP1 (26–28). Our observation that PARP1- selective inhibition and PARP1/2 inhibition have similar effects on proliferation and AR-driven transcription support findings in other disease settings that PARP1-selective inhibition have comparable or improved efficacy with lower hematologic toxicities (29–32).

Previous studies reported that AR directly regulates expression of DNA damage repair genes (1, 3, 17), which sensitized HR-proficient prostate cancer to PARP inhibition (2). However, the lack of obvious synergy between PARP inhibition and AR inhibition in clinical studies with HR-proficient patients suggests that this model of AR regulated DNA repair may not be as generalizable as previously thought (4–6). We were surprised to find that androgen deprivation did not regulate DNA repair in our studies, although this supports our later finding that AR inhibition and PARP inhibition do not synergize in HR-proficient models. While in contrast to the accepted model of AR-regulated DNA repair, others have also observed that AR does not directly regulate expression of DNA damage response genes (33, 34).

This work raises several key points for clinical use of PARP inhibitors in prostate cancer. While our results do not support the hypothesis that AR inhibition and PARP inhibition synergize in CRPC, this does not preclude the possibility that some patients will experience clinical benefit from the independent additive effect of these two treatments. It is also important to note that these results are likely not generalizable to HR-deficient prostate cancer. In the context of HR-deficiency, the combination of AR inhibition and PARP inhibition dramatically prolongs progression-free survival (4–6). However, since these tumors are typically sensitive to both PARP inhibition and AR inhibition as single agents, the efficacy of the drug combination may be due to additive effects of inhibiting two distinct pathways with non-overlapping mechanisms of resistance. It remains unclear whether there are additional synergistic effects between PARP inhibition and AR inhibition in HR-deficient prostate cancer.

Additionally, a key finding of this study is that inhibiting cell cycle progression impairs response to PARP inhibitors, suggesting that PARP inhibitors should not be combined with agents that induce cell cycle arrest. For these experiments, we used CDK4/6 inhibition with palbociclib to induce G1 arrest (35, 36). CDK4/6 inhibition is not widely used in prostate cancer, but many other common treatments, including androgen deprivation, also induce G1 arrest (37, 38). This effect does not seem to be specific to PARP inhibitors, but rather a widely generalizable effect of G1 arrest on response to DNA damaging agents (Fig. 5E-H and ref. (39, 40). Interestingly, patients with HR-deficiency respond poorly to androgen deprivation therapy compared to patients with HR-proficient tumors (41, 42). This suggests that part of the improved efficacy of the AR inhibition and PARP inhibitor combination treatment in this disease setting may be due to a decreased cell cycle arrest upon androgen deprivation treatment.

## METHODS

### Cell Culture

LNCaP and 22RV1 were maintained in RPMI-1640 (Gibco) supplemented with 10% fetal bovine serum (FBS) (Sigma-Aldrich). LNCaP-abl and LN95 cells were maintained in phenol red-free RPMI-1640 (Gibco) supplemented with 10% charcoal dextran treated FBS (Omega Scientific). Cells were authenticated by STR genotyping (ATCC) and verified to be mycoplasma free (Mycoplasma Detection Kit, Invivogen).

### Cell Viability Assays

Cells were seeded in 96-well plates, allowed to adhere overnight, and treated in quadruplicate with olaparib (Selleck Chemicals), rucaparib (Selleck Chemicals), AZD-5305 (MedChemExpress), enzalutamide (Selleck Chemicals), palbociclib (Selleck Chemicals), cisplatin (Fresenius Kabi), etoposide (Accord Healthcare), or doxorubicin (Sigma Aldrich) as indicated for 7 d. Cell viability was measured using the Cell Titer Glo Luminescent Cell Viability Assay (Promega) according the manufacturer’s instructions. Treatments were normalized to their respective controls.

### DNA Damage Assays

Cells were plated on glass coverslips and treated as indicated. Cells were then fixed in 1% methanol-free formaldehyde in PBS, permeabilized in 0.1% Triton X-100 in PBS, and blocked for 1 h in 1% BSA in PBS. Cells were incubated with primary antibody (phospho-histone H2AX_S139_; Cell Signaling Technology #9718, 1:300 dilution) for 1 h at room temperature, followed by secondary antibody incubation (donkey anti-rabbit IgG Alexa-Fluor488, Invitrogen # A32790 1:2000 dilution) for 1 h at room temperature. Coverslips were mounted in ProLong Glass Antifade Mountant with NucBlue Stain (Invitrogen). Slides were imaged at 63X using a Zeiss 980 Airyscan confocal microscope. Foci were counted in >100 cells per treatment group by manual counting by an investigator blinded to treatment group.

### RNA-sequencing

Cells were treated in duplicate with PARP inhibition (5 μM) and/or DHT (1 nM). RNA was extracted using the RNeasy Mini Kit (Qiagen) with on-column DNase I digestion according to the manufacturer’s instructions. Libraries were prepared and sequenced by Novogene with Illumina Novaseq6000 paired-end 150 bp sequencing. The VIPER pipeline (43) was used for STAR alignment to the hg19 genome (44), read count normalization using Cufflinks (45) quality control with RSeQC (46) and differential expression analysis using DESeq2 (47). Genes with a log2-fold change > |1| and p-adjusted >0.05 were considered differentially expressed. The data generated in this study are available within this article and its supplementary files (Table S1).

### ATAC-sequencing

Cells were treated in duplicate with olaparib (5 μM) and/or DHT (1 nM). Nuclei were harvested from 100,000 cells per replicate. Samples and libraries were prepared using the Active Motif ATAC-seq kit according to the manufacturer’s instructions. Samples were sequenced on the Illumina NovaSeq X platform with paired-end 150 bp reads, with 50 million reads/sample. The CoBRA pipeline (48) was used for sample alignment to the hg19 genome and identification of differential peaks using DESeq2 (47). Peaks with a log2-fold change > |1| and p-adjusted >0.05 were considered differentially expressed. The data generated in this study are publicly available in this article and it’s supplementary files (Table S2).

### Immunoblotting

Cells were lysed in RIPA buffer (Boston Bioproducts) with protease inhibitor cocktail (Sigma-Aldrich). Concentrations were measured and normalized using the Pierce BCA Protein Quantification Assay kit (ThermoFisher Scientific) according to the manufacturer’s instructions. Samples were prepared for immunoblot in LDS sample buffer (Invitrogen) and heated to 95°C for 5 min to denature protein. Proteins were resolved on a NuPage 4-12% Bis-Tris gel (ThermoFisher Scientific) and transferred to nitrocellulose membranes using the Trans-Blot Turbo system (BioRad). Transfer was confirmed by Ponceau S. Membranes were blocked in 5% milk powder in tris-buffered saline with 0.1% Tween-20 (TBST), then incubated overnight in primary antibody overnight at 4°C (AR, Santa-Cruz #7305, 1:1000 dilution; GAPDH, Cell Signaling Technology #2118, 1:1000 dilution). Membranes were blotted in HRP-conjugated secondary antibody (Goat anti-mouse IgG, Invitrogen #31430, 1:5000 dilution) for 1 h at room temperature. Blots were developed in Western Blotting Luminol Reagent (Santa Cruz Biotechnology) and imaged with the ChemiDoc system (BioRad).

## ACKNOWLEDGEMENTS

This work was supported by the DoD PCRP Early Investigator Research Award HT94252310910 to N.A.T and NIH 5P01CA163227 to M.B. This work utilized an Illumina NovaSeq X Plus that was purchased with funding from a National Institutes of Health SIG grant 1S10OD036228-01.

## Notes

**Conflicts of Interest Disclosure Statement:** MB received sponsored research support and was formerly a consultant to and SAB member for Novartis. MB formerly served on the SAB of FibroGen. MB serves on the SAB of Kronos Bio and holds equity in the company. MB serves on the SAB of GV20 Therapeutics and holds equity in the company. EC served as a paid consultant to DotQuant, and received Institutional sponsored research funding unrelated to this work from Astra Zeneca, AbbVie, Gilead, Sanofi, Zenith Epigenetics, Bayer Pharmaceuticals, Forma Therapeutics, Genentech, GSK, Janssen Research, Kronos Bio, Foghorn Therapeutics, K36, and MacroGenics. All other authors declare no potential conflicts of interest.

### Competing Interest Statement

MB received sponsored research support and was formerly a consultant to and SAB member for Novartis. MB formerly served on the SAB of FibroGen. MB serves on the SAB of Kronos Bio and holds equity in the company. MB serves on the SAB of GV20 Therapeutics and holds equity in the company. EC served as a paid consultant to DotQuant, and received Institutional sponsored research funding unrelated to this work from Astra Zeneca, AbbVie, Gilead, Sanofi, Zenith Epigenetics, Bayer Pharmaceuticals, Forma Therapeutics, Genentech, GSK, Janssen Research, Kronos Bio, Foghorn Therapeutics, K36, and MacroGenics. All other authors declare no potential conflicts of interest.

## REFERENCES

1. Goodwin JF, Schiewer MJ, Dean JL, Schrecengost RS, de Leeuw R, Han S, et al. A hormone-DNA repair circuit governs the response to genotoxic insult. Cancer Discov. 2013;3(11):1254–71.

2. Li L, Karanika S, Yang G, Wang J, Park S, Broom BM, et al. Androgen receptor inhibitor-induced “BRCAness” and PARP inhibition are synthetically lethal for castration-resistant prostate cancer. Sci Signal. 2017;10(480).

3. Jividen K, Kedzierska KZ, Yang CS, Szlachta K, Ratan A, Paschal BM. Genomic analysis of DNA repair genes and androgen signaling in prostate cancer. BMC Cancer. 2018;18(1):960.

4. Saad F, Clarke NW, Oya M, Shore N, Procopio G, Guedes JD, et al. Olaparib plus abiraterone versus placebo plus abiraterone in metastatic castration-resistant prostate cancer (PROpel): final prespecified overall survival results of a randomised, double-blind, phase 3 trial. Lancet Oncol. 2023;24(10):1094–108.

5. Fizazi K, Azad AA, Matsubara N, Carles J, Fay AP, De Giorgi U, et al. First-line talazoparib with enzalutamide in HRR-deficient metastatic castration-resistant prostate cancer: the phase 3 TALAPRO-2 trial. Nat Med. 2024;30(1):257–64.

6. Chi KN, Sandhu S, Smith MR, Attard G, Saad M, Olmos D, et al. Niraparib plus abiraterone acetate with prednisone in patients with metastatic castration-resistant prostate cancer and homologous recombination repair gene alterations: second interim analysis of the randomized phase III MAGNITUDE trial. Ann Oncol. 2023;34(9):772–82.

7. Cobb J. ODAC marathon: Committee charges through a pileup of clinical questions in a two-day, four-application session. The Cancer Letter. 2025;51:6–27.

8. Schreiber V, Ame JC, Dolle P, Schultz I, Rinaldi B, Fraulob V, et al. Poly(ADP-ribose) polymerase-2 (PARP-2) is required for efficient base excision DNA repair in association with PARP-1 and XRCC1. J Biol Chem. 2002;277(25):23028–36.

9. Pinnola A, Naumova N, Shah M, Tulin AV. Nucleosomal core histones mediate dynamic regulation of poly(ADP-ribose) polymerase 1 protein binding to chromatin and induction of its enzymatic activity. J Biol Chem. 2007;282(44):32511–9.

10. Krishnakumar R, Gamble MJ, Frizzell KM, Berrocal JG, Kininis M, Kraus WL. Reciprocal binding of PARP-1 and histone H1 at promoters specifies transcriptional outcomes. Science. 2008;319(5864):819–21.

11. Frizzell KM, Gamble MJ, Berrocal JG, Zhang T, Krishnakumar R, Cen Y, et al. Global analysis of transcriptional regulation by poly(ADP-ribose) polymerase-1 and poly(ADP-ribose) glycohydrolase in MCF-7 human breast cancer cells. J Biol Chem. 2009;284(49):33926–38.

12. Krishnakumar R, Kraus WL. PARP-1 regulates chromatin structure and transcription through a KDM5B-dependent pathway. Mol Cell. 2010;39(5):736–49.

13. Gui B, Gui F, Takai T, Feng C, Bai X, Fazli L, et al. Selective targeting of PARP-2 inhibits androgen receptor signaling and prostate cancer growth through disruption of FOXA1 function. Proc Natl Acad Sci U S A. 2019;116(29):14573–82.

14. Ju BG, Lunyak VV, Perissi V, Garcia-Bassets I, Rose DW, Glass CK, et al. A topoisomerase IIbeta-mediated dsDNA break required for regulated transcription. Science. 2006;312(5781):1798–802.

15. Ju BG, Solum D, Song EJ, Lee KJ, Rose DW, Glass CK, et al. Activating the PARP-1 sensor component of the groucho/ TLE1 corepressor complex mediates a CaMKinase IIdelta-dependent neurogenic gene activation pathway. Cell. 2004;119(6):815–29.

16. Pavri R, Lewis B, Kim TK, Dilworth FJ, Erdjument-Bromage H, Tempst P, et al. PARP-1 determines specificity in a retinoid signaling pathway via direct modulation of mediator. Mol Cell. 2005;18(1):83–96.

17. Polkinghorn WR, Parker JS, Lee MX, Kass EM, Spratt DE, Iaquinta PJ, et al. Androgen receptor signaling regulates DNA repair in prostate cancers. Cancer Discov. 2013;3(11):1245–53.

18. Tran EU, Royz E, Yamamoto K, Marley S, Song A, Pan E, et al. Bipolar androgen therapy for treatment of metastatic castration-resistant prostate cancer: A case series. Prostate. 2025;85(1):40–7.

19. Schweizer MT, Antonarakis ES, Wang H, Ajiboye AS, Spitz A, Cao H, et al. Effect of bipolar androgen therapy for asymptomatic men with castration-resistant prostate cancer: results from a pilot clinical study. Sci Transl Med. 2015;7(269):269ra2.

20. Denmeade SR, Wang H, Agarwal N, Smith DC, Schweizer MT, Stein MN, et al. TRANSFORMER: A Randomized Phase II Study Comparing Bipolar Androgen Therapy Versus Enzalutamide in Asymptomatic Men With Castration-Resistant Metastatic Prostate Cancer. J Clin Oncol. 2021;39(12):1371–82.

21. Teply BA, Wang H, Luber B, Sullivan R, Rifkind I, Bruns A, et al. Bipolar androgen therapy in men with metastatic castration-resistant prostate cancer after progression on enzalutamide: an open-label, phase 2, multicohort study. Lancet Oncol. 2018;19(1):76–86.

22. Chatterjee P, Schweizer MT, Lucas JM, Coleman I, Nyquist MD, Frank SB, et al. Supraphysiological androgens suppress prostate cancer growth through androgen receptor-mediated DNA damage. J Clin Invest. 2019;129(10):4245–60.

23. Schweizer MT, Gulati R, Yezefski T, Cheng HH, Mostaghel E, Haffner MC, et al. Bipolar androgen therapy plus olaparib in men with metastatic castration-resistant prostate cancer. Prostate Cancer Prostatic Dis. 2023;26(1):194–200.

24. Traphagen NA, Schwartz GN, Tau S, Roberts AM, Jiang A, Hosford SR, et al. Estrogen Therapy Induces Receptor-Dependent DNA Damage Enhanced by PARP Inhibition in ER+ Breast Cancer. Clin Cancer Res. 2023;29(18):3717–28.

25. Rajan P, Sudbery IM, Villasevil ME, Mui E, Fleming J, Davis M, et al. Next-generation sequencing of advanced prostate cancer treated with androgen-deprivation therapy. Eur Urol. 2014;66(1):32–9.

26. Johannes JW, Balazs A, Barratt D, Bista M, Chuba MD, Cosulich S, et al. Discovery of 5-4-[(7-Ethyl-6-oxo-5,6-dihydro-1,5-naphthyridin-3-yl)methyl]piperazin-1-yl-N-methylpyridine-2-carboxamide (AZD5305): A PARP1-DNA Trapper with High Selectivity for PARP1 over PARP2 and Other PARPs. J Med Chem. 2021;64(19):14498–512.

27. Gao S, Hou Y, Xu Y, Li J, Zhang C, Jiang S, et al. Discovery of Pyrazolo[1,5,4-de]quinoxalin-2(3H)-one Derivatives as Highly Potent and Selective PARP1 Inhibitors. J Med Chem. 2024;67(23):21380–99.

28. Johannes JW, Balazs AYS, Barratt D, Bista M, Chuba MD, Cosulich S, et al. Discovery of 6-Fluoro-5-4-[(5-fluoro-2-methyl-3-oxo-3,4-dihydroquinoxalin-6-yl)methyl]piperazin-1-yl-N-methylpyridine-2-carboxamide (AZD9574): A CNS-Penetrant, PARP1-Selective Inhibitor. J Med Chem. 2024;67(24):21717–28.

29. Illuzzi G, Staniszewska AD, Gill SJ, Pike A, McWilliams L, Critchlow SE, et al. Preclinical Characterization of AZD5305, A Next-Generation, Highly Selective PARP1 Inhibitor and Trapper. Clin Cancer Res. 2022;28(21):4724–36.

30. Dellavedova G, Decio A, Formenti L, Albertella MR, Wilson J, Staniszewska AD, et al. The PARP1 Inhibitor AZD5305 Impairs Ovarian Adenocarcinoma Progression and Visceral Metastases in Patient-derived Xenografts Alone and in Combination with Carboplatin. Cancer Res Commun. 2023;3(3):489–500.

31. Staniszewska AD, Pilger D, Gill SJ, Jamal K, Bohin N, Guzzetti S, et al. Preclinical Characterization of AZD9574, a Blood-Brain Barrier Penetrant Inhibitor of PARP1. Clin Cancer Res. 2024;30(7):1338–51.

32. Dall G, Vandenberg CJ, Nesic K, Ratnayake G, Zhu W, Vissers JHA, et al. Targeting homologous recombination deficiency in uterine leiomyosarcoma. J Exp Clin Cancer Res. 2023;42(1):112.

33. Hasterok S, Scott TG, Roller DG, Spencer A, Dutta AB, Sathyan KM, et al. The Androgen Receptor Does Not Directly Regulate the Transcription of DNA Damage Response Genes. Mol Cancer Res. 2023;21(12):1329–41.

34. Gao S, Gao Y, He HH, Han D, Han W, Avery A, et al. Androgen Receptor Tumor Suppressor Function Is Mediated by Recruitment of Retinoblastoma Protein. Cell Rep. 2016;17(4):966–76.

35. Fry DW, Harvey PJ, Keller PR, Elliott WL, Meade M, Trachet E, et al. Specific inhibition of cyclin-dependent kinase 4/6 by PD 0332991 and associated antitumor activity in human tumor xenografts. Mol Cancer Ther. 2004;3(11):1427–38.

36. Toogood PL, Harvey PJ, Repine JT, Sheehan DJ, VanderWel SN, Zhou H, et al. Discovery of a potent and selective inhibitor of cyclin-dependent kinase 4/6. J Med Chem. 2005;48(7):2388–406.

37. Knudsen KE, Arden KC, Cavenee WK. Multiple G1 regulatory elements control the androgen-dependent proliferation of prostatic carcinoma cells. J Biol Chem. 1998;273(32):20213–22.

38. Agus DB, Cordon-Cardo C, Fox W, Drobnjak M, Koff A, Golde DW, et al. Prostate cancer cell cycle regulators: response to androgen withdrawal and development of androgen independence. J Natl Cancer Inst. 1999;91(21):1869–76.

39. Dean JL, McClendon AK, Knudsen ES. Modification of the DNA damage response by therapeutic CDK4/6 inhibition. J Biol Chem. 2012;287(34):29075–87.

40. Pikman Y, Alexe G, Roti G, Conway AS, Furman A, Lee ES, et al. Synergistic Drug Combinations with a CDK4/6 Inhibitor in T-cell Acute Lymphoblastic Leukemia. Clin Cancer Res. 2017;23(4):1012–24.

41. Fettke H, Dai C, Kwan EM, Zheng T, Du P, Ng N, et al. BRCA-deficient metastatic prostate cancer has an adverse prognosis and distinct genomic phenotype. EBioMedicine. 2023;95:104738.

42. Annala M, Struss WJ, Warner EW, Beja K, Vandekerkhove G, Wong A, et al. Treatment Outcomes and Tumor Loss of Heterozygosity in Germline DNA Repair-deficient Prostate Cancer. Eur Urol. 2017;72(1):34–42.

43. Cornwell M, Vangala M, Taing L, Herbert Z, Koster J, Li B, et al. VIPER: Visualization Pipeline for RNA-seq, a Snakemake workflow for efficient and complete RNA-seq analysis. BMC Bioinformatics. 2018;19(1):135.

44. Dobin A, Davis CA, Schlesinger F, Drenkow J, Zaleski C, Jha S, et al. STAR: ultrafast universal RNA-seq aligner. Bioinformatics. 2013;29(1):15–21.

45. Trapnell C, Williams BA, Pertea G, Mortazavi A, Kwan G, van Baren MJ, et al. Transcript assembly and quantification by RNA-Seq reveals unannotated transcripts and isoform switching during cell differentiation. Nat Biotechnol. 2010;28(5):511–5.

46. Wang L, Wang S, Li W. RSeQC: quality control of RNA-seq experiments. Bioinformatics. 2012;28(16):2184–5.

47. Love MI, Huber W, Anders S. Moderated estimation of fold change and dispersion for RNA-seq data with DESeq2. Genome Biol. 2014;15(12):550.

48. Qiu X, Feit AS, Feiglin A, Xie Y, Kesten N, Taing L, et al. CoBRA: Containerized Bioinformatics Workflow for Reproducible ChIP/ATAC-seq Analysis. Genomics Proteomics Bioinformatics. 2021;19(4):652–61.

