## Supplemental Figures for "Lack of synergy between AR targeted therapies and PARP inhibitors in homologous recombination-proficient prostate cancer"

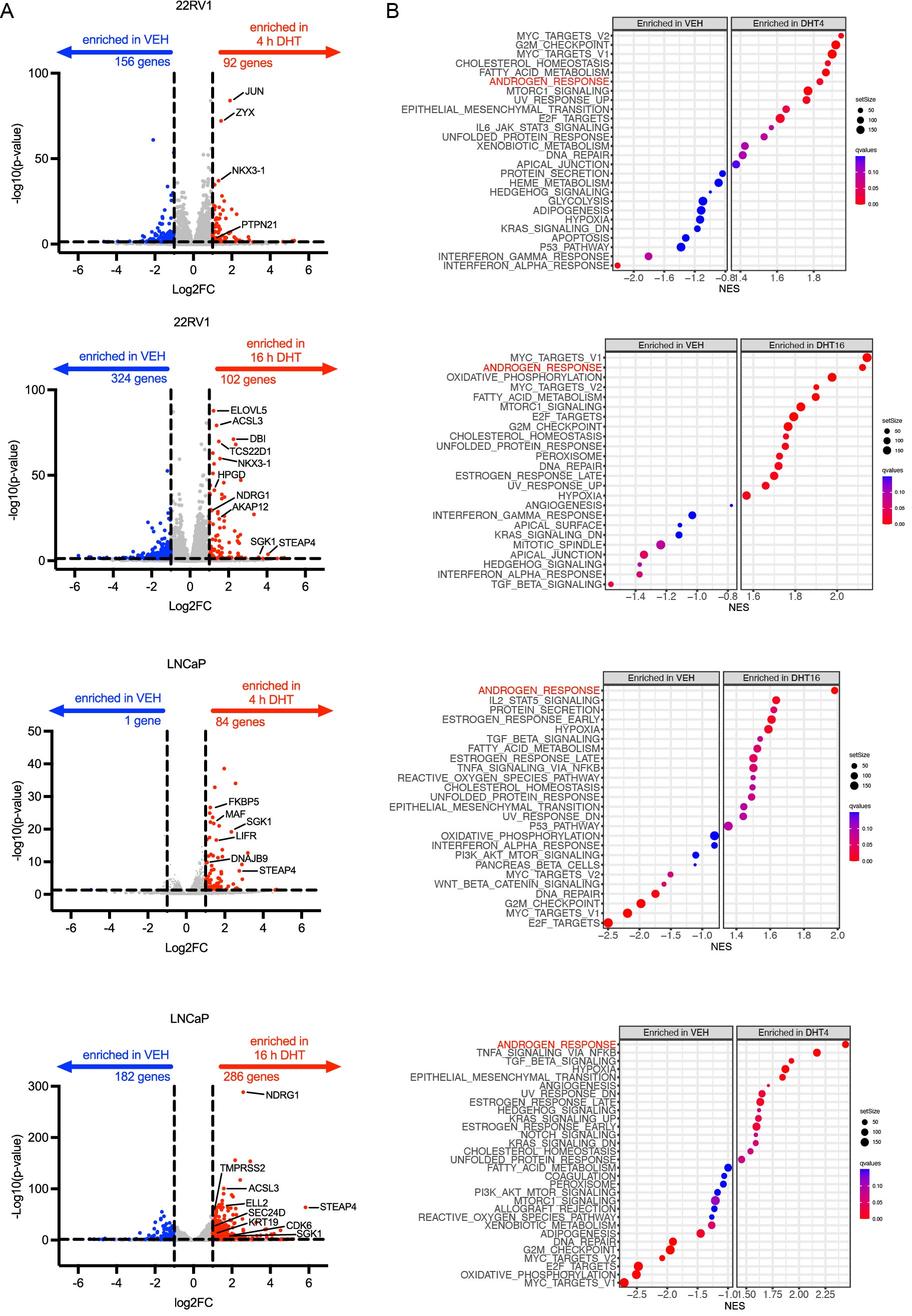
**Figure S1. DHT treatment induces a robust androgen response signature.** (A) Genes with significantly differential expression (Log2FC > |1|; p-adjusted <0.05) at 4 h or 16 h of treatment with DHT compared to vehicle (androgen-deprived) treated cells. (B) Normalized enrichment scores for Hallmarks pathways in DHT treated vs. vehicle treated cells.

**Figure S2. Differentially expressed genes in LNCaP upon PARPi treatment.** Genes with significantly differential expression (Log2FC > |1|; p-adjusted <0.05) following treatment ± PARPi in vehicle (androgen-deprived) media or upon DHT stimulation in LNCaP cells

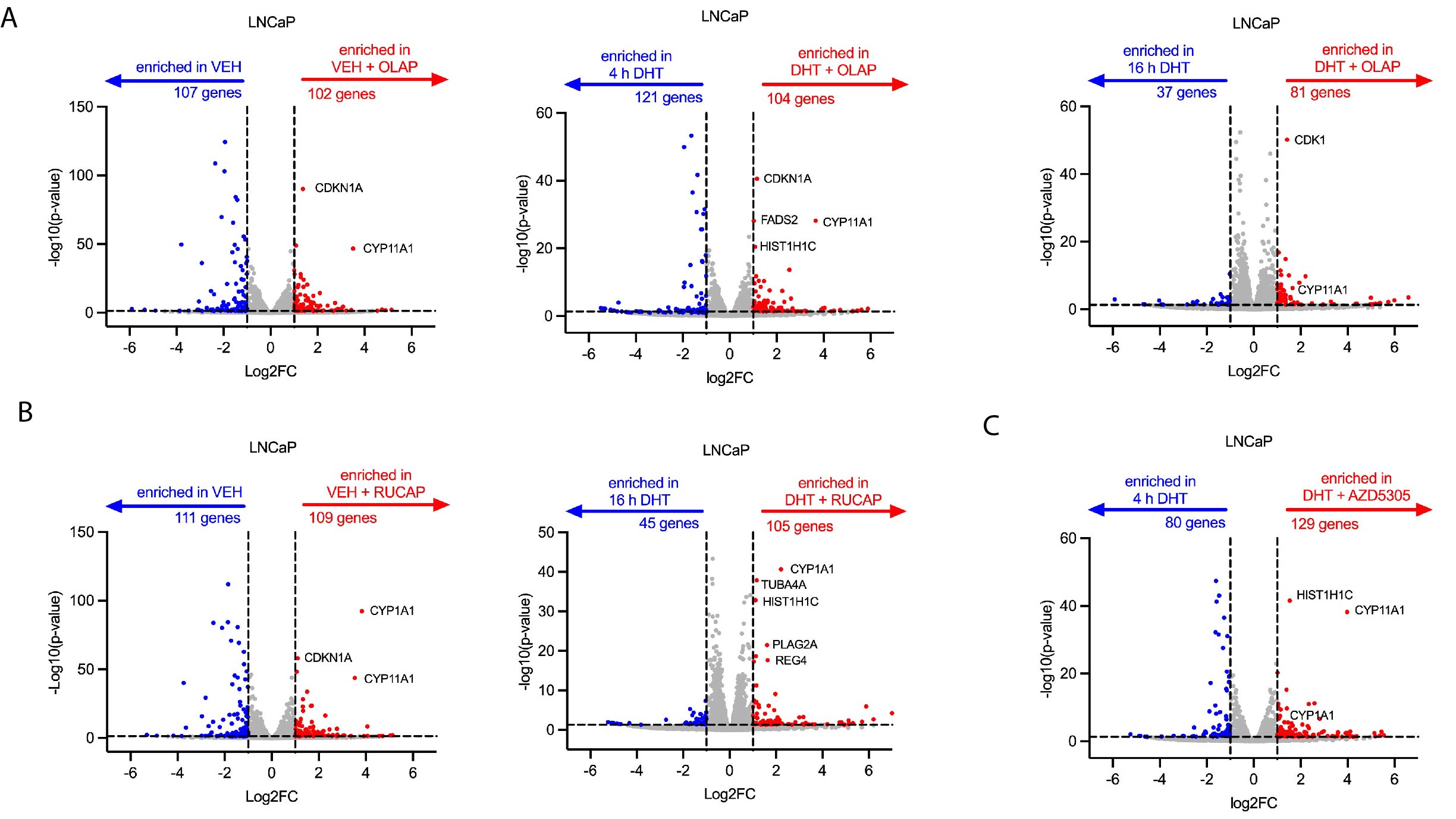

**Figure S3. Differentially expressed genes in 22RV1 upon PARPi treatment.** Genes with significantly differential expression (Log2FC > |1|; p-adjusted <0.05) following treatment ± PARPi in vehicle (androgen-deprived) media or upon DHT stimulation in 22RV1 cells.

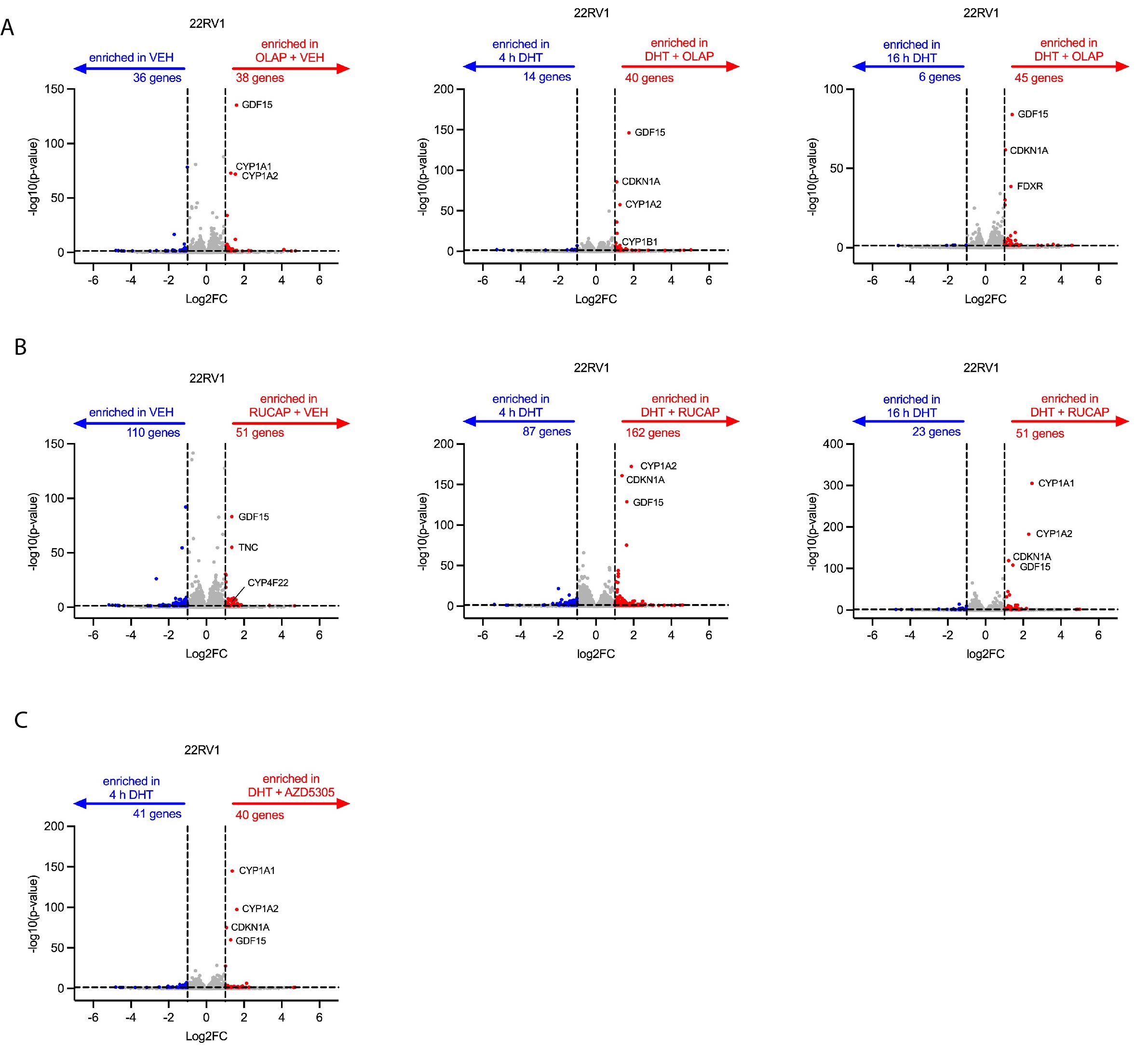

**Figure S4. Hallmarks pathway enrichment for AR and PARPi regulated gene clusters.** Enrichment of Hallmarks pathways for each androgen-regulated gene cluster in (A) 22RV1 and (B) LNCaP cells. Dotted line shows significance threshold of p = 0.05.

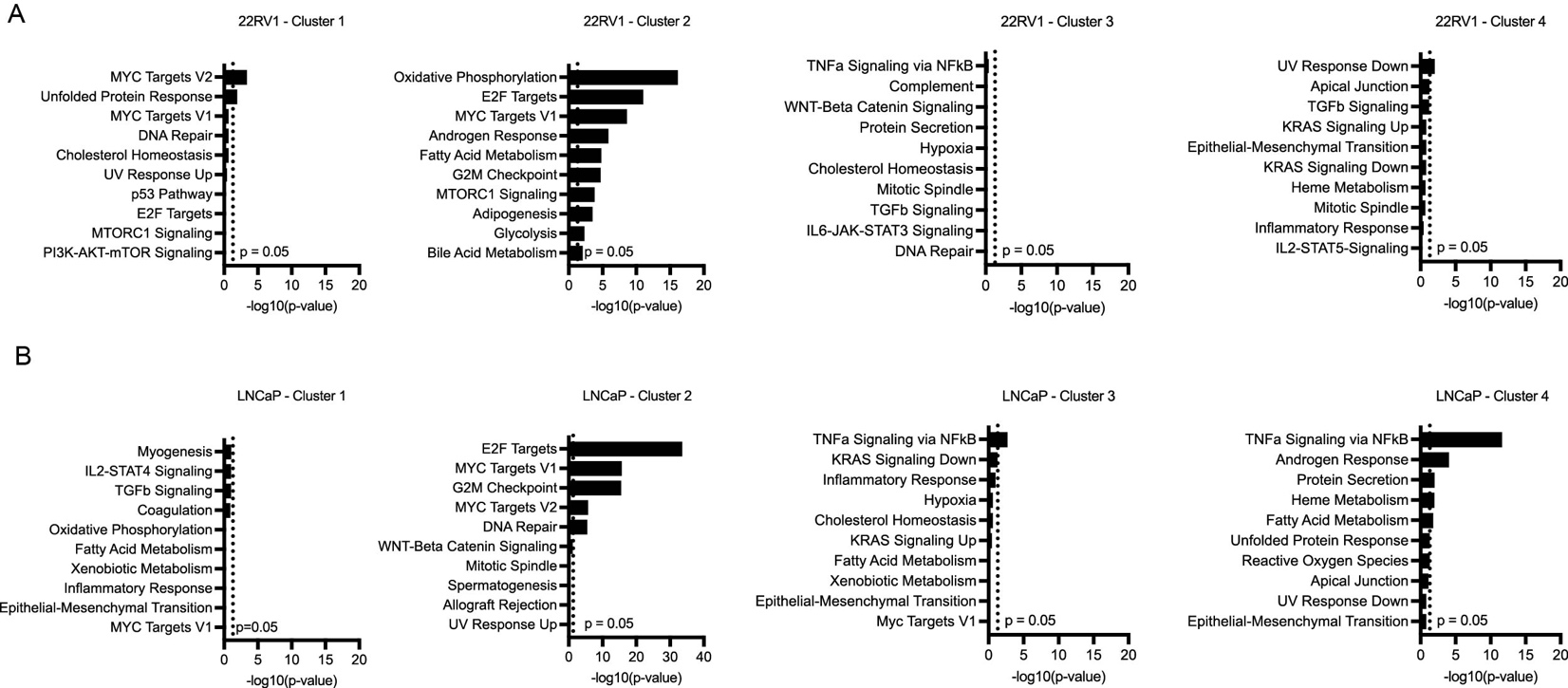

**Figure S5. DHT treatment induces robust changes in chromatin accessibility.** (A) Chromatin regions with significantly differential peaks (Log2FC > |1|; p-adjusted <0.05) upon 24h treatment with DHT in 22RV1 and LNCaP cells. (B) ATAC-seq peaks *for LRRC41*, a Cluster 2 AR-regulated gene in 22RV1. Expression of this gene is induced by DHT but decreased upon treatment with PARPi.

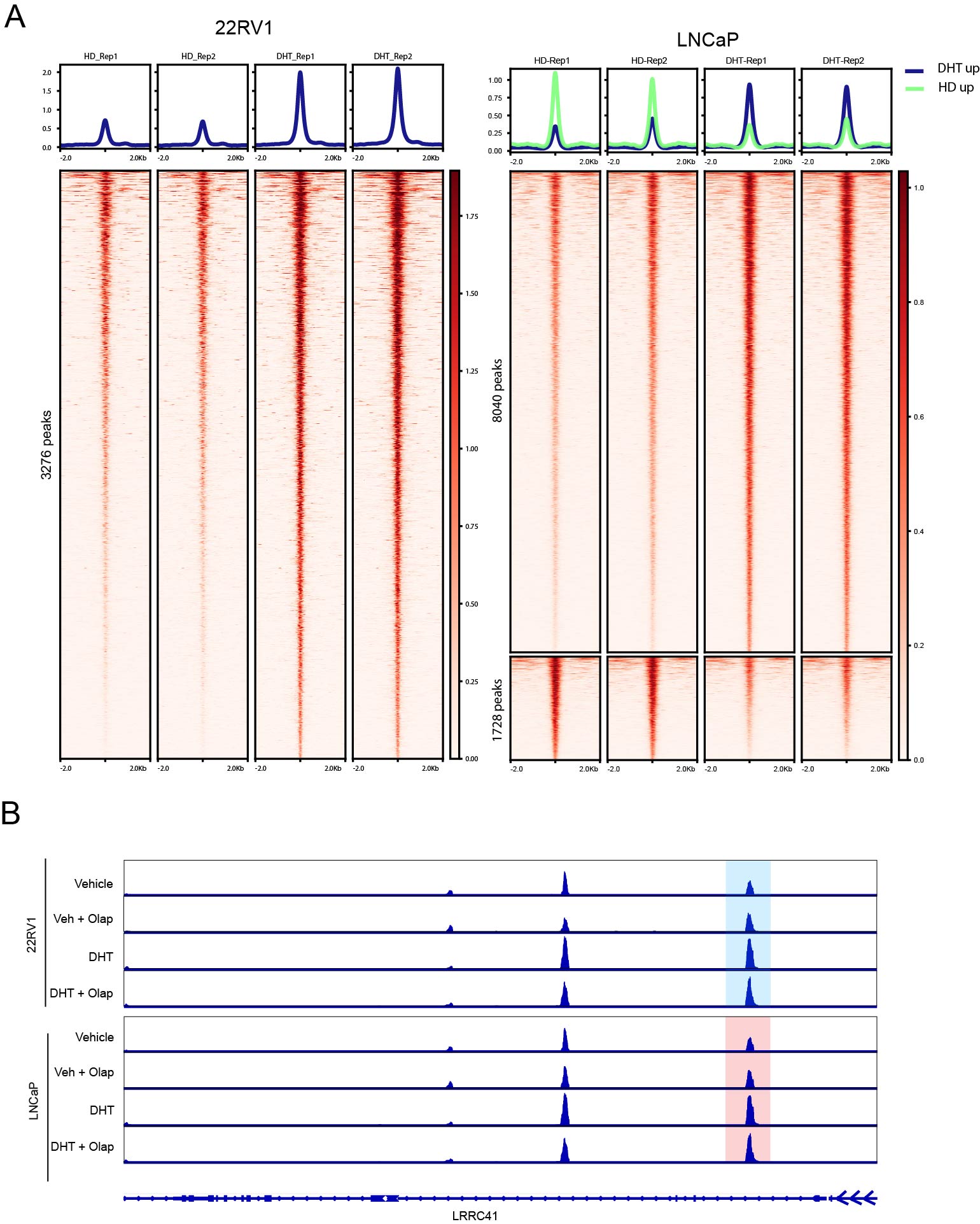

**Figure S6.**

**Treatment with enzalutamide prevents LNCaP cells from acquiring DNA damage upon PARPi treatment.** LNCaP cells were seeded in full media and treated ± 10 μM enzalutamide (ENZA), ± 5 μM olaparib as indicated for 72 h. Cells were fixed and stained for γH2AX (green), with NucBlue (blue) nuclear counterstaining. γH2AX foci were quantified in >100 cells per treatment group. ***p<0.0005, n.s. = not significant by one-way ANOVA with Bonferroni correction for multiple comparisons.

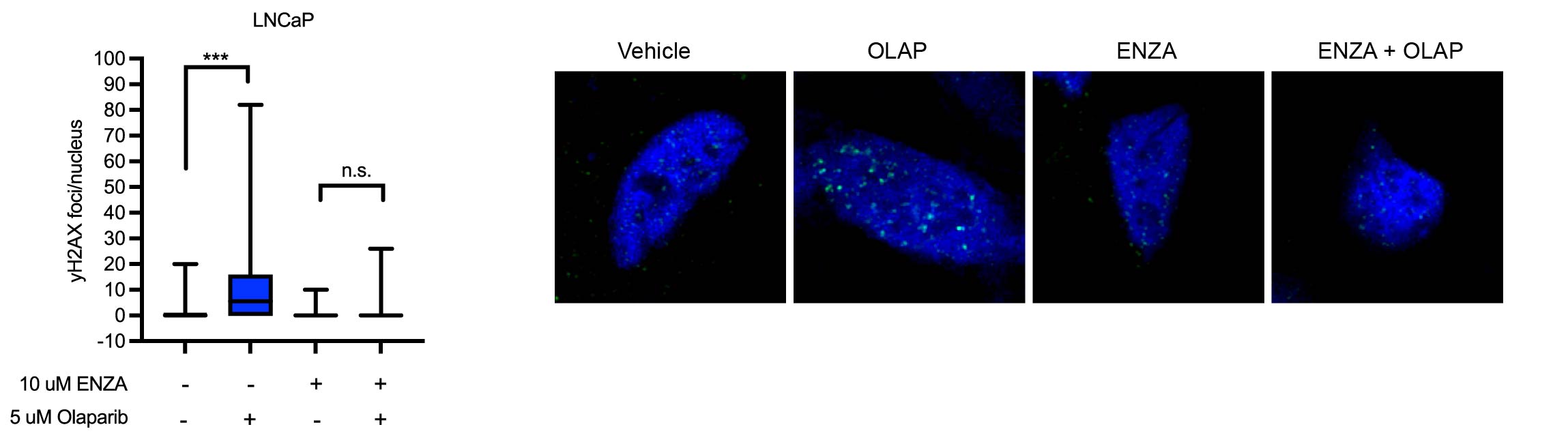

**Figure S7. Cell cycle inhibition impairs growth and therapeutic response to PARPi.** (A) LNCaP cells were seeded in full media and treated with palbociclib in quadruplicate as indicated for 7 d prior to assessing cell viability. (B) CRPC cells were seeded in growth media and treated ± 5 μM palbociclib in quadruplicate for 7 d prior to assessing cell viability. (C) LNCaP cells were seeded in full media ± 5 μM palbociclib and treated with PARPi as indicated in quadruplicate for 7 d prior to assessing cell viability. Data are shown as mean ± SD. ***p<0.0005 by Student’s two-tailed t-test.

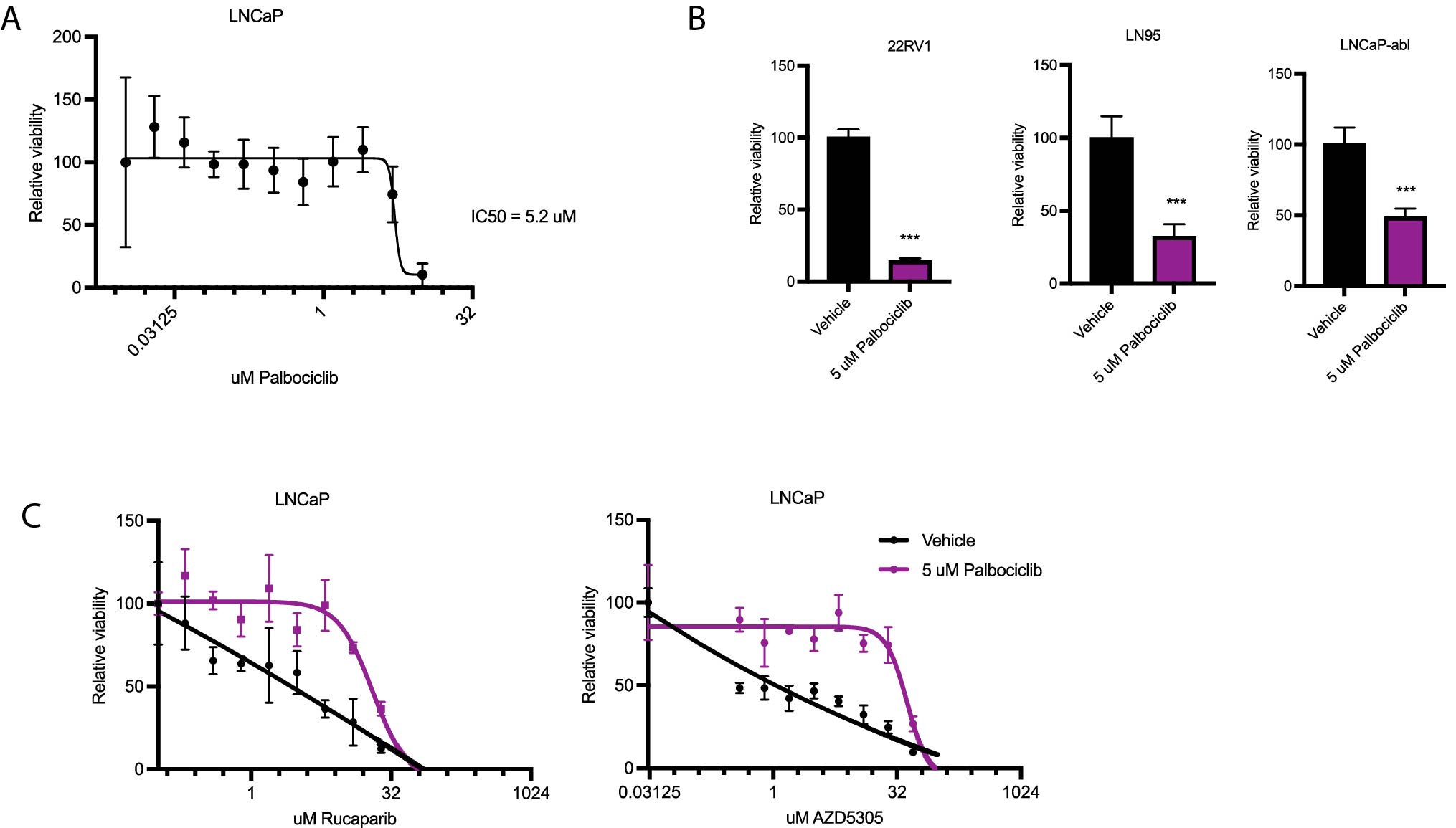

**Figure S8. Cell cycle inhibition impairs therapeutic response to DNA-damaging chemotherapies.** (A) Cells were seeded in androgen-deprived media ± 1 nM DHT and treated with cisplatin as indicated in quadruplicate prior to assessing cell viability. (B) Cells were seeded in growth media ± 10 μM enzalutamide (ENZA) and treated with cisplatin as indicated for 7 d prior to assessing cell viability. (C) LNCaP cells were seeded in full media ± 5 μM palbociclib, then treated with doxorubicin or etoposide as indicated for 7 d prior to assessing cell viability. (D) CRPC cells were seeded in growth media ± 5 μM palbociclib, then treated with cisplatin as indicated for 7 d prior to assessing cell viability. Data are shown as mean ± SD.

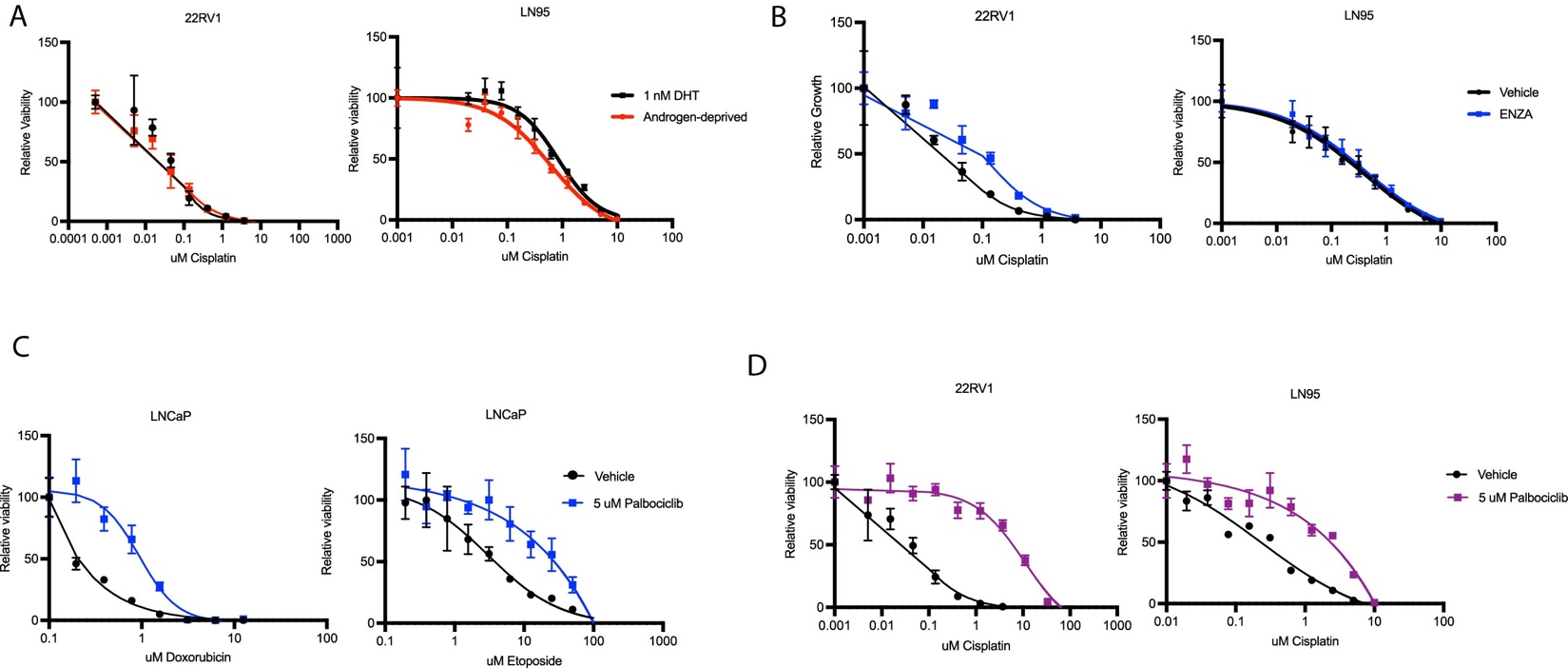
